# Inhibition of FOXM1 synergizes with BCL2 inhibitor Venetoclax in killing non-t(11;14) multiple myeloma cells via repressing MYC pathway

**DOI:** 10.1101/2024.09.27.613548

**Authors:** Zhi Wen, Siegfried Janz, Yidan Wang, Benita S. Katzenellenbogen, John A. Katzenellenbogen, Sung Hoon Kim, Adedayo Onitilo

## Abstract

Despite significant improvements in the prognosis of Multiple Myeloma (MM), relapsed/refractory MM remains a major challenge. BCL2 inhibitor Venetoclax induced complete or very good partial responses in 6% of non-t(11;14) MM cases, compared to 27% in t(11;14) cases, when used as monotherapy in relapsed/refractory MM. Though Venetoclax was proposed to treat t(11;14) cases, the resistance became a concern. Furthermore, non-t(11;14) cases account for 80-85% of MM cases, which underscores the value of Venetoclax in non-t(11;14) MM. Here, we report a recently-invented small molecule inhibitor of FOXM1 NB73 synergizing with Venetoclax in killing MM cells. FOXM1, a critical forkhead box transcription factor in high-risk and relapsed/refractory MM, represents a promising therapeutic target of MM. We examined the mechanisms underlying the synergies of Venetoclax and NB73 using multi-omics and molecular and cellular biology tools in non-t(11;14) myeloma cell lines with high FOXM1 expression. NB73 induces immediate loss of FOXM1, decreases BCL2 expression, and increases Puma expression in myeloma cells. Venetoclax enhances NB73-induced FOXM1 ubiquitination and degradation. The NB73-Venetoclax combination abrogates the binding of FOXM1 to the promoters of genes in the MYC pathway, such as PLK1, MYC, CDC20, and CCNA2, leading to the repression of the transcription of these MYC pathway genes. The PLK1-specific inhibitor GSK461364 synergies with NB73 in suppressing myeloma cell growth. Therefore, NB73 synergizes with Venetoclax in killing myeloma cells. Conclusively, the NB73-Venetoclax combination abolishes FOXM1-mediated transcriptional activation of the MYC pathway, resulting in intensive apoptosis of myeloma cells without t(11;14) but with high FOXM1 expression.

**Statement of significance:** This study implicates that targeting FOXM1 will alleviate resistance to BCL2 inhibitor Venetoclax in non-t(11;14) myeloma cells expressing high FOXM1.

## Introduction

With an annual incidence of more than 30,000 cases, Multiple Myeloma (MM) ranks as the second most common blood cancer in the United States [1]. Owing to recently developed myeloma drugs and newly FDA-approved immunotherapies, the announcement of a 5-year relative survival rate of 61% in 2023 marks a significant improvement in MM prognosis compared to 52% reported in 2020 [1]. However, relapsed and refractory myeloma remains a dominant obstacle to further improving survival rates. Concerningly, relapse cases from Car-T cell and bispecific antibody therapies have emerged as significant challenges in MM, with continuous observations of relapse in recent years [2].

The BCL2 family consists of anti-apoptotic, pro-apoptotic, and BH3-domain-only subfamilies [3]. BCL2 is anti-apoptotic, BAK is pro-apoptotic, and Puma belongs to the BH3-domain-only subfamily. BCL2 binds to BAK and prevents BAK from releasing cytochrome c, thereby halting cell apoptosis. However, Puma displaces BAK from the BCL2-BAK complex to initiate apoptosis [4]. Venetoclax, the first FDA-approved BH3 mimetic inhibiting BCL2 [5], is a potent apoptosis inducer used to treat chronic lymphocytic leukemia and newly-diagnosed acute myeloid leukemia. It has also been extensively evaluated in clinical trials, such as the CANOVA study, for treating a subgroup of MM patients with t(11;14) translocations [6, 7]. Recent studies have revealed that t(11;14) MM cells can be either Venetoclax-sensitive or Venetoclax-resistant depending on their expression pattern of B-cell genes [8–10]. Moreover, Venetoclax has induced a complete response or very good partial response in 6% of non-t(11;14) MM cases, compared to 27% in t(11;14) cases, in the M13-367 clinical trial using Venetoclax monotherapy in relapsed/refractory MM patients [11]. As non-t(11;14) MM cases comprise 80-85% of all MM cases, Venetoclax may hold significance in non-t(11;14) MM patients, such as these with hyperactivated BCL2 pathway [7]. Though non-t(11;14) MM cell lines appear resistant to Venetoclax [10], the underlying mechanisms remain unclear.

FOXM1 is a transcription factor in the forkhead box family, which governs many aspects of malignant growth and is widely acknowledged as a master gene in both hematopoietic and solid cancers [12]. It has been implicated in newly diagnosed high-risk myeloma [13], relapsed myeloma [14], and the development of drug resistance in myeloma [15, 16]. In addition to promoting myeloma cell cycle progression, FOXM1 acts as a promoter of the bioenergetic pathways of glycolysis and oxidative phosphorylation, providing fuel for MM cells [17]. Deletion of FOXM1 alleles significantly suppresses MM cell growth in both in vitro and in vivo settings [17]. Alternatively, NB73 is a small-molecule inhibitor of FOXM1 based on a 1,1-diarylethylene scaffold, which binds to and reduces FOXM1 protein [18]. NB73, on its own and in combination with an HSP90 inhibitor, has demonstrated the ability to suppress breast cancer and MM both in vitro and in vivo [17, 18]. Furthermore, NB73 has exhibited favorable safety and pharmacokinetic profiles in experimental animals, underscoring its potential as a promising candidate for clinical trials and combinatorial treatments [19].

Considering the broad involvement of FOXM1 in newly diagnosed high-risk and relapsed/refractory MM, we designed drug combinations of NB73 and FDA-approved anti-cancer drugs, testing their synergies in non-t(11;14) MM cell lines with high FOXM1 expression. Herein, we have addressed the synergy of NB73 and Venetoclax in non-t(11;14) MM cell lines with high FOXM1 expression, which provides the mechanism perspective to the recent publication of the killing efficacy of xenograft in NSG mice and primary myeloma cells [20].

## Materials and Methods

### ZIP drug synergy scoring assay

The drugs were diluted sequentially in Opti-MEM medium (GIBCO, cat# 31985070) in 8-stripe PCR tubes. Subsequently, 5 µL of each diluted drug was aliquoted into each well of a 96-well plate using an 8-channel P20 pipetman (Gilson, cat# F144070). Next, 100 µL of 0.3×10^6^ cells/mL Delta47/OPM2 cells were seeded into each well using an 8-channel P200 pipetman (Gilson, cat# F144072). The cells were then cultured at 5% CO2, 37°C for two days. Afterward, 20 µL of CellTiter-Glo cell viability substrate (Promega, cat# G7573) was added to each well using a DISTRIMAN Repetitive Pipette (Gilson, cat# F164001) and incubated at room temperature for 10 minutes on a microplate shaker (USA Scientific, cat# 7402-4000). Finally, luminescence was measured using a BMG LUMIstar Omega plate reader. The cell viability in the DMSO control was set as 100% for normalization. Synergy between drugs was assessed using the ZIP synergy score, as per instructions at https://synergyfinder.fimm.fi. A ZIP Average Score > 10 indicates synergy between drugs, a ZIP Average Score < -10 indicates antagonism, while a ZIP Average Score between -10 and 10 suggests an additive effect.

### Chromatin Immunoprecipitation (ChIP)

14 mL of OPM2 or Delta47 cells at a concentration of 0.3 × 10^6^ cells/mL were evenly distributed into 10-cm dishes. After 24 hours, the following treatments were administered: DMSO, NB73 (2 µM), Venetoclax (5 µM), and the NB73-Venetoclax combination for OPM2 cells, while DMSO, NB73 (1.6 µM), Venetoclax (5 µM), and the NB73-Venetoclax combination for Delta47 cells. After another 24 hours of incubation, cells were collected for ChIP using the anti-FOXM1 antibody (Cell signaling, cat# 20459) and the Pierce Magnetic ChIP Kit (Thermo, cat# 26157). The ChIP procedure followed the manufacturer’s instructions, including sonication of the cell nuclei for two cycles of 15 seconds pulse and 1 minute rest at 20% energy using a Branson SFX250 sonicator, addition of 2 µg antibody per ChIP, and washing of the beads-immunoprecipitates four times. Genomic DNA (gDNA) was then extracted for subsequent qPCR and next-generation deep sequencing analyses.

All other materials and methods *can be found in supplemental files or previously published* [21–23].

## Results

### FOXM1 inhibitor NB73 synergizes with BCL2 inhibitor Venetoclax in killing myeloma cells

Myeloma OPM2 and Delta47 cells express significantly higher levels of FOXM1 compared to the other nine myeloma cell lines [17]. Suppression of FOXM1 using either its small-molecule inhibitor NB73 or the CRISPR/Cas9 knockout tool induced growth inhibition in both OPM2 and Delta47 cells in vitro and in vivo [17], highlighting the crucial role of FOXM1 in cell survival. To assess the drug synergy, we conducted in vitro assays combining NB73 with anti-MM drugs. Daratumumab, Dexamethasone, Car T-cells, and bispecific antibodies leverage the MM patient’s immune system to target MM cells, an aspect not replicated in traditional in vitro single-cell-population culture systems. Therefore, we combined NB73 with one of six anti-MM drugs (Bortezomib, Cyclophosphamide, Lenalidomide, Selinexor, Vincristine, and Venetoclax) in the in vitro drug tests in OPM2 cells (Fig. 1A). The ZIP drug synergy assay (Average score > 10) was employed to evaluate drug synergy [24, 25]. Notably, Venetoclax demonstrated synergy with NB73 in suppressing both OPM2 and Delta47 cells (Fig. 1B-C), suggesting that inhibiting FOXM1 enhances the cell-killing effects of the BCL2 inhibitor in FOXM1-high MM cells. As BCL2 is a key anti-apoptotic protein, we selected one dose combination with a high ZIP score and measured cell apoptosis one day after drug treatments. Figure 1B illustrated that Venetoclax minimally induced cell apoptosis, consistent with its known efficacy in treating t(11;14) MM, which OPM2 cells lack. In contrast, NB73 induced substantial cell apoptosis in OPM2 cells, a response markedly enhanced by Venetoclax (Fig. 1B). This augmentation of cell apoptosis was similarly observed in Delta47 cells (Fig. 1C). Finally, this synergy was also observed in NSG mice engrafted with OPM2 cells as well as in ex vivo assay with a new one-day culturing protocol [20].

**Figure 1:**
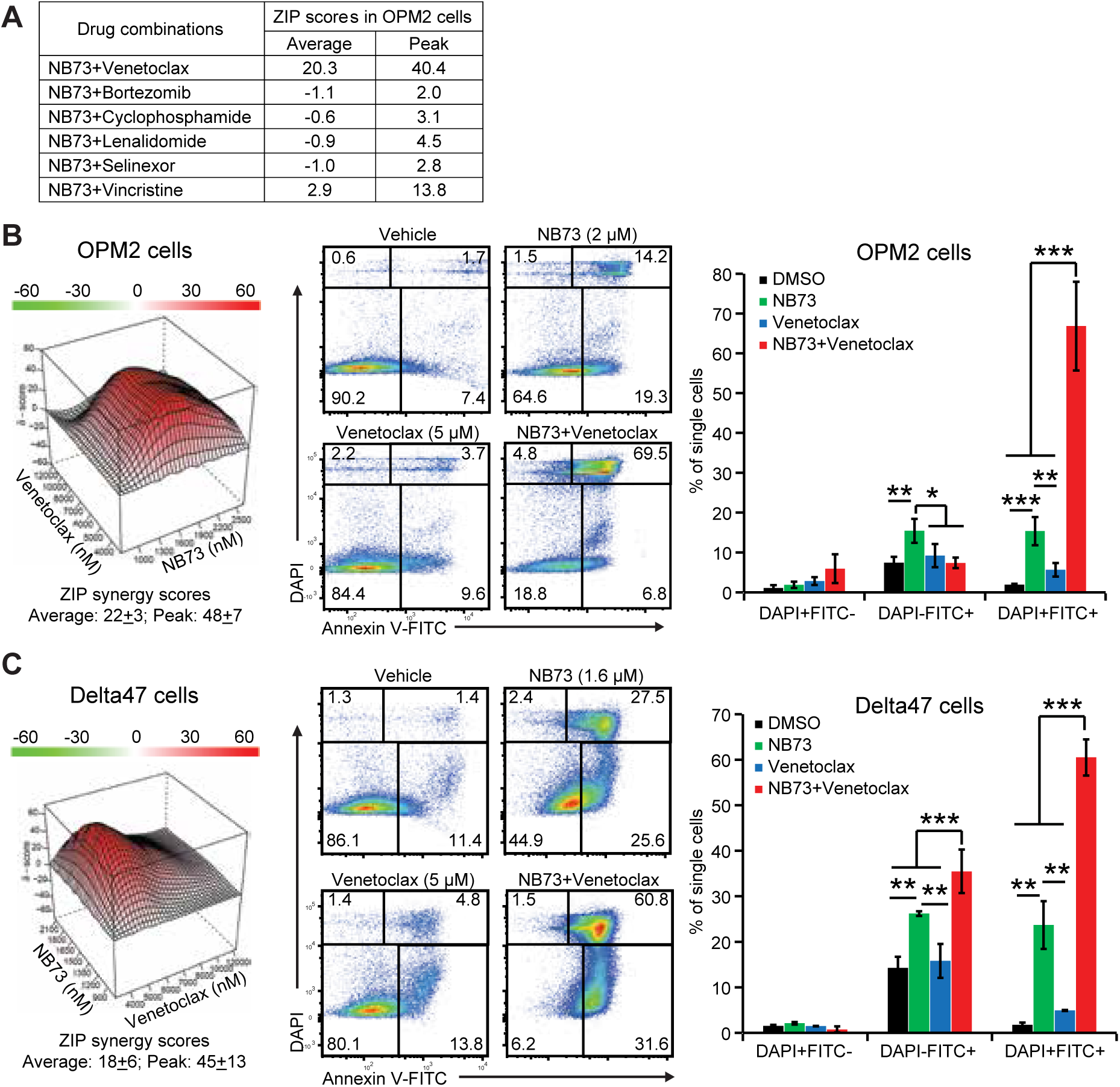
BCL2 inhibitor Venetoclax synergizes with FOXM1 inhibitor NB73 in suppressing myeloma cells. OPM2 and Delta47 cells, exhibiting the highest FOXM1 levels among 11 myeloma cell lines [17], were treated with the specified drugs for 2 days. Cell viabilities were assessed using CellTiter-Glo assay and normalized to DMSO controls. The cell apoptosis assays were conducted 1 day after treatment. (A) Average and Peak ZIP drug synergy scores of NB73 and 6 FDA-approved anti-MM drugs in OPM2 cells were calculated using the online tool at www.synergyfinder.fimm.fi. (B-C) Venetoclax exhibited synergy with NB73 in inducing cell apoptosis in OPM2 cells (B) and Delta47 cells (C). *left*: ZIP drug synergy scoring assays (8×9 matrix) of NB73 and Venetoclax treatments in myeloma cells, depicted as average ± SD of triplicates (n=3); *middle*: representative flow images of Annexin-V binding assays in myeloma cells treated with NB73 and/or Venetoclax; *right*: histogram of Annexin-V binding assays (n=3). *p* values were calculated by Student’s t-test with two tails. *: *p*<0.05; **: *p*<0.01; ***: *p*<0.001.

FOXM1^-/-^ cells exhibited significantly slower growth rates compared to FOXM1^+/+^ cells [17]. Transcriptomic analysis suggested a down-regulation of the BCL2 pathway in FOXM1^-/-^ OPM2 cells (supplemental Fig. S1A). Interestingly, Venetoclax decreased BCL2 RNA level and increased Puma RNA level in FOXM1^-/-^ cells compared to DMSO (supplemental Fig. S1B), implicating FOXM1 in Venetoclax bioactivities in OPM2 cells. Dose-response efficacy studies revealed an enhanced sensitivity to Venetoclax in FOXM1^-/-^ OPM2 cells (supplemental Fig. S1C), supporting that FOXM1 promotes resistance to Venetoclax in OPM2 cells. Conversely, deleting FOXM1 did not markedly down-regulate the BCL2 pathway in Delta47 cells (supplemental Fig. S1D). Actually, Venetoclax increased RNA levels of both BCL2 and Puma in FOXM1^-/-^ Delta47 cells, introducing a conflict to the BCL2 pathway (supplemental Fig. S1E). The dose-response efficacy studies of FOXM1^-/-^ and FOXM1^+/+^ Delta47 cells showed a moderate improvement in sensitivity to Venetoclax in FOXM1^-/-^ cells (supplemental Fig. S1F). Overall, we identified that NB73 synergizes with Venetoclax in killing myeloma cells in vitro.

### Unlike FOXM1 knockout, NB73 induces transcriptomic changes of immediate FOXM1 loss

Deletion of FOXM1 alleles leads to marked growth inhibition of OPM2 and Delta47 cells, but the stable cell lines have managed to survive and continue proliferating [17]. Transcriptomic studies of FOXM1^-/-^ OPM2 cells revealed changes in the stable state. For instance, we observed up-regulations of the MYC and E2F pathways, which promote cell proliferation (supplemental Table-S1). There may be differences in the transcriptome between stable loss of FOXM1 in FOXM1^-/-^ cells and immediate loss of FOXM1 induced by NB73. Therefore, we conducted an RNA-seq assay to explore transcriptomic changes in OPM2 cells treated with DMSO and NB73, respectively. NB73 up-regulated 3,055 genes (fold change > 2 and p < 0.05), including tumor-suppressing genes such as p53, IFIT3, and TNF, among others (Fig. 2A-B). Interestingly, NB73 strongly increased the expression of BMP2, Bone Morphogenetic Protein-2 (Fig. 2A). BMP2 is a growth factor secreted by osteoblasts that promotes bone formation. FDA has approved BMP2 as an osteoinductive growth factor to serve as a bone graft substitute for inducing bone regeneration [26]. By stimulating BMP2 expression, NB73 may antagonize the activated osteoclasts in MM patients, thereby improving myeloma bone disease. In contrast, NB73 down-regulated 2,275 genes, including growth-promoting genes such as CCNE1 and DKK1, as well as the glycolysis-related enzyme FBP2 (Fig. 2A).

**Figure 2:**
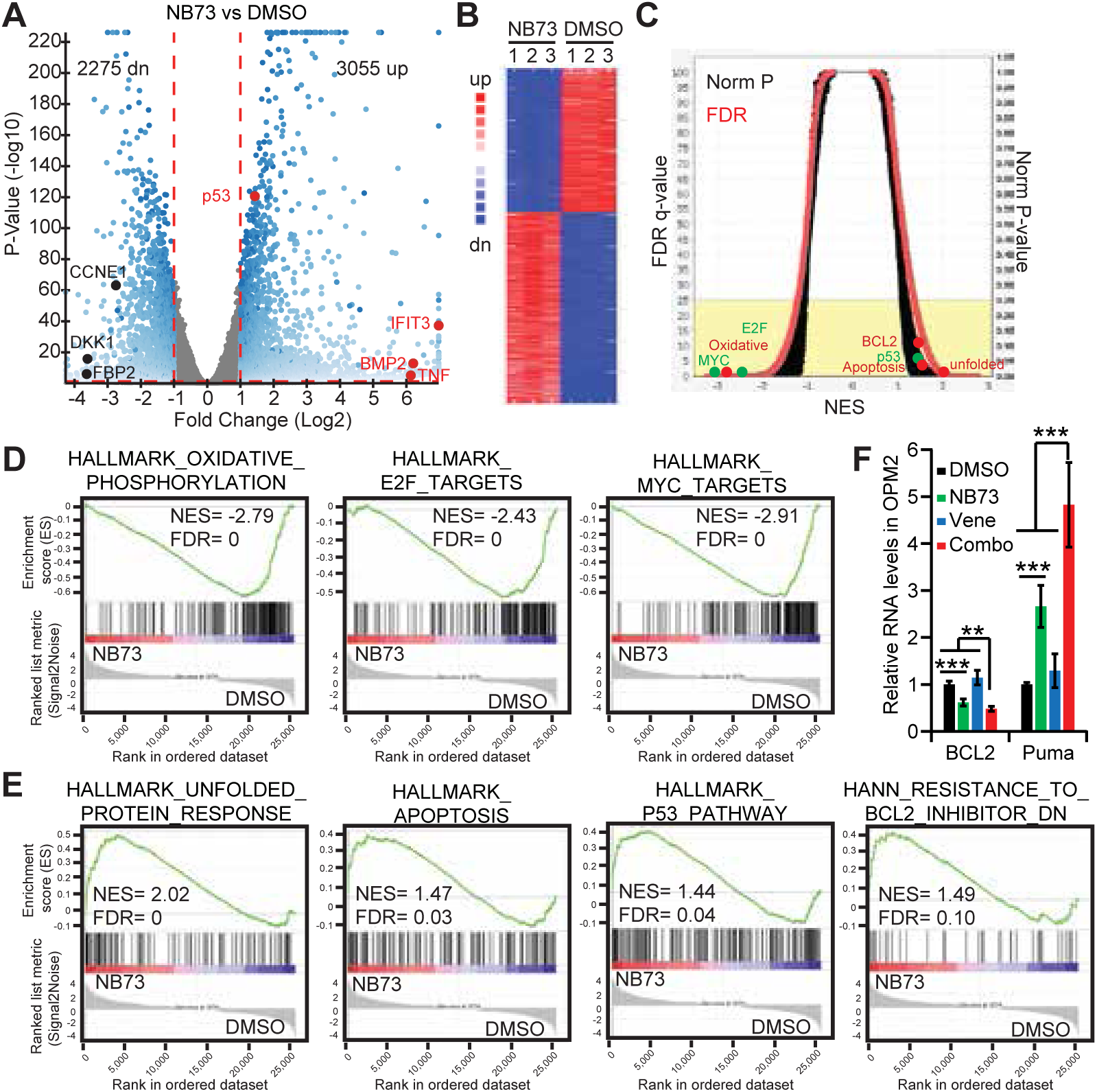
Decipher transcriptomic changes induced by NB73 in OPM2 cells. OPM2 cells were treated with NB73 or DMSO for 24 hours. RNA was extracted for RNA-seq study. (A) Volcano plot representing all transcripts, with red dots indicating growth-suppressing genes and black dots representing growth-promoting genes. (B) Heat map depicting the expression profiles of the most differentially expressed genes between NB73-treated and control groups in OPM2 cells. (C) Overview of Gene Set Enrichment Analysis (GSEA) results, highlighting significant pathway alterations induced by NB73 treatment. (D) GSEA showing enrichment of pathways related to oxidative phosphorylation, MYC and E2F pathways in the control group. (E) GSEA showing enrichment of pathways associated with unfolded protein response (UPR), p53 signaling, apoptosis, and BCL2 inhibitor-related pathways in the NB73 group. (F) RNA levels of BCL2 and Puma were measured with qRT-PCR assays in FOXM1^+/+^ OPM2 cells treated with DMSO, NB73, Venetoclax and the combination, respectively. *p* values were calculated by Student’s t-test with two tails. **: *p*<0.01; ***: *p*<0.001.

GSEA studies revealed substantial pathway changes (Fig. 2C). For example, NB73 significantly inhibited growth-promoting pathways, including MYC, E2F, and oxidative phosphorylation (Fig. 2D), whereas they were up-regulated in FOXM1^-/-^ OPM2 cells (supplemental Table-S1). Additionally, we observed up-regulations of tumor-suppressing pathways (e.g., apoptosis and p53) and the myeloma-specific unfolded protein response (UPR) pathway (Fig. 2E). Myeloma cells are characterized by high endogenous UPR due to overproduction of immunoglobulins, which can be further stimulated by exogenous ER stresses from proteasome inhibitors, leading to myeloma cell apoptosis [27]. Furthermore, NB73 up-regulated a set of genes that were down-regulated in BCL2 inhibitor-resistant cells (Fig. 2E), which was supported by qRT-PCR assays showing the decreased BCL2 RNA levels and increased Puma RNA levels induced by both NB73 and the NB73-Venetoclax combination (Fig. 2F). This observation implicated that NB73 disrupts the BCL2 pathway, facilitating the cell apoptosis. Unlike in FOXM1^-/-^ OPM2 cells (supplemental Fig. S1B), Venetoclax alone did not regulate the RNA levels of BCL2 and Puma RNA in FOXM1^+/+^ OPM2 cells (Fig. 2F), emphasizing the roles of FOXM1 in the resistance to Venetoclax.

Indeed, the comparative analysis of GSEA results between immediate FOXM1 loss induced by NB73 and stable FOXM1 loss in FOXM1^-/-^ cells revealed distinct transcriptomic patterns (supplemental Table-S1). These differences may suggest unique mechanisms underlying myeloma cell survival following FOXM1-tageted treatment. The transcriptomic changes observed in FOXM1^-/-^ cells could recapitulate long-term effects of FOXM1 inhibition, therefore shedding light on potential mechanisms of resistance to FOXM1 inhibition.

### The NB73-Venetoclax combination represses MM cells through three synergy modes

To dissect the molecular mechanisms underlying the synergy, we treated OPM2 cells with DMSO, NB73, Venetoclax, and the combination, respectively, for RNA-seq assays. As depicted in Figure 3A-B, the cluster of Venetoclax was very close to DMSO, suggesting mild transcriptomic changes between these two groups. This result correlated with the slight difference in cell apoptosis between DMSO and Venetoclax (Fig. 1B). In contrast, there were notable transcriptomic changes between DMSO, NB73, and the combination (Fig. 3A-B). Compared to DMSO, the combination up-regulated 3,513 genes (fold change > 2 and p < 0.05) while down-regulating 2,597 genes (Fig. 3C), which was substantially higher than the changes observed in NB73 vs DMSO (Fig. 2A). Among these genes, we observed that the combination enhanced the NB73-induced up-regulation of tumor-suppressing genes such as XAF1, TNF, AIM2, TNFSF15, and IFIT3 (Fig. 3D), demonstrating the synergistic effects between NB73 and Venetoclax.

**Figure 3:**
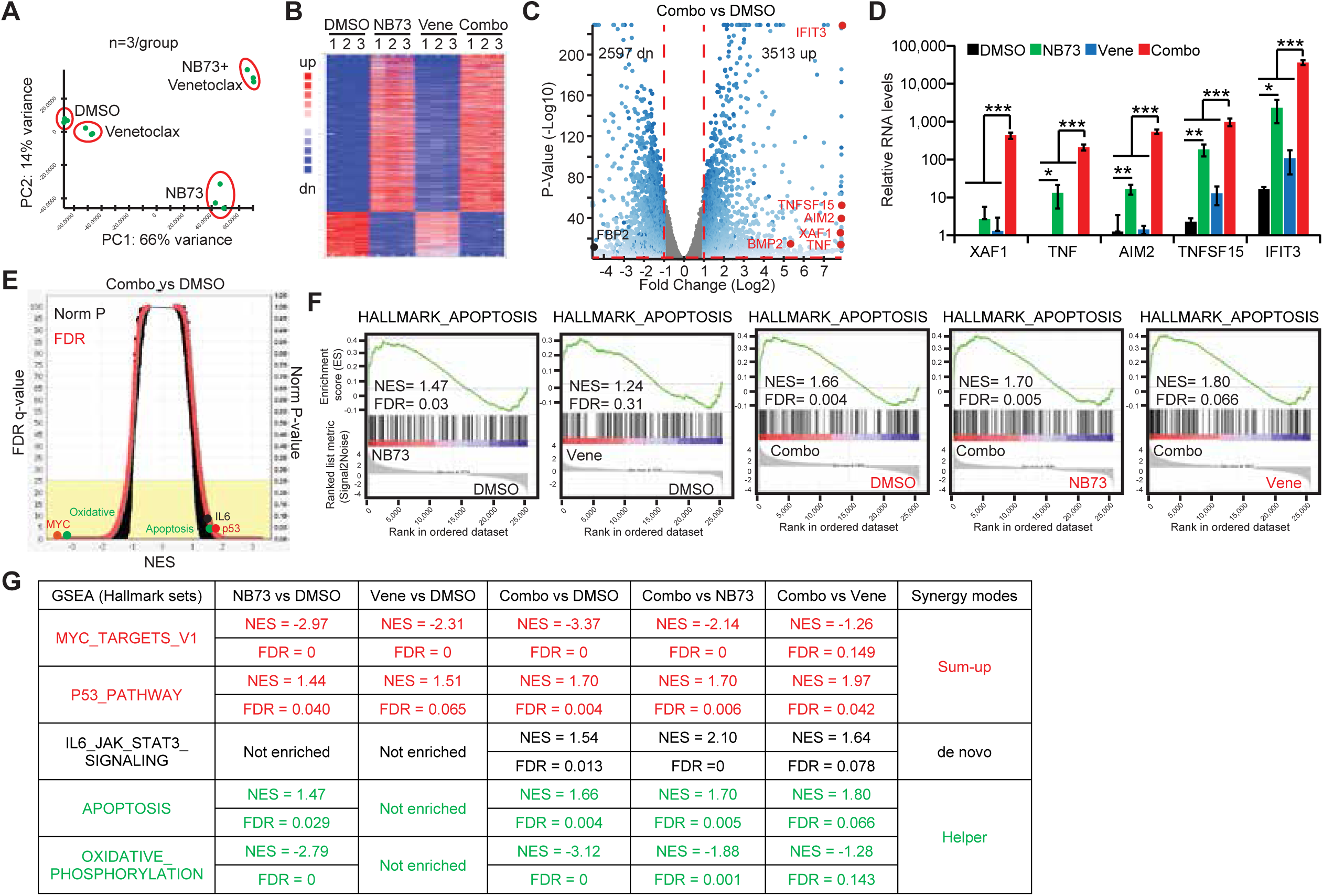
Transcriptomics study reveals drug synergy modes in OPM2 cells. OPM2 cells were treated with DMSO, NB73, Venetoclax, or the combination for 24 hours, respectively. RNA was then extracted for RNA-seq analysis. (A) Principal component analysis (PCA) of 12 samples demonstrates distinct clustering based on treatment conditions. (B) Heatmap depicts the expression levels of the most differentially expressed genes in the combo vs. DMSO group, highlighting treatment-induced transcriptional changes. (C) Volcano plot illustrates differential gene expression between the combo and DMSO groups, with red dots indicating growth-suppressing genes and black dots representing growth-promoting genes. (D) Representative RNA levels of growth-suppressing genes are shown to elucidate treatment effects. (E) Overview of GSEA results reveals pathway alterations induced by treatment. (F) Multiple comparisons of the Apoptosis pathway among DMSO, NB73, Venetoclax, or the combo groups show the drug synergy. (G) Drug synergy modes are visualized using the GSEA tool, with red indicating sum-up mode, black representing de novo mode, and green indicating helper mode.

The GSEA study between DMSO and the combination treatment displayed remarkable changes in many important tumor-associated pathways (Fig. 3E). We utilized the GSEA tool to conduct multiple comparisons, including NB73 vs DMSO, Venetoclax vs DMSO, the combination vs DMSO, the combination vs NB73, and the combination vs Venetoclax (Fig. 3F-G). For example, both NB73 and the combination enriched the apoptosis pathway when compared to DMSO, whereas Venetoclax did not (Fig. 3F). Moreover, the combination further enriched the apoptosis pathway compared to NB73 or Venetoclax alone, suggesting that the combination is a stronger inducer of apoptosis than NB73 alone (Fig. 3F). Such multiple comparisons not only identified the stronger player in a specific pathway but also elucidated the modes of synergies (Fig. 3G). We observed three modes of synergies: (1) sum-up mode, in which both drugs enriched a gene set and the combination was stronger than each single drug (e.g., MYC pathway); (2) de novo mode, in which neither drug enriched a gene set, but the combination gained enrichment (e.g., IL6 pathway); and (3) helper mode, in which only one drug enriched a gene set while the combination became stronger than the single drug (e.g., Apoptosis pathway). In summary, the transcriptomic studies provided valuable insights into the mechanisms underlying drug synergy, such as inspecting the MYC pathway.

### The NB73-Venetoclax combination abolishes MYC signaling in myeloma cells

Hyperactivation of MYC pathway is a key driver of carcinogenesis prevalent in MM [28–30]. We also reported a mouse model of advanced MM by introducing *Nras^Q61R^* to *Vκ-MYC* mice [31]. The inhibition of MYC pathway by NB73 was significantly enhanced by the addition of Venetoclax (Fig. 3G). We focused on four critical genes in MYC pathway, including MYC, PLK1, CDC20, and CCNA2, which were among the top down-regulated genes (Fig. 4A). These genes are known to promote growth, and their inhibition induces cell growth inhibition and apoptosis. We first validated the NB73-mediated down-regulation of these four genes at the RNA level using qRT-PCR and at the protein level using immunoblotting in OPM2 cells (Fig. 4B-C). Venetoclax further enhanced the NB73-mediated down-regulation of these genes in OPM2 cells (Fig. 4B-C). Next, we treated OPM2 cells with a PLK1-specific inhibitor, GSK461364 [32], alone and in combination with NB73, and observed an average ZIP synergy score over 10 (Fig. 4D), indicating synergy between the FOXM1 inhibitor and PLK1 inhibitor. As GSK461364 arrests cell cycle at G2-M phase [32], we selected one dose combination with a high ZIP score and measured cell cycle one day after drug treatments. Though GSK461364 arrested OPM2 cells at G2-M phase, the combination did not enhance this arrest (Fig. 4E-F). Similarly, NB73 induced OPM2 cell apoptosis while the combination did not enhance the cell apoptosis either (Fig. 4G-H). Actually, the NB73-GSK461364 combination not only induced cell apoptosis but also arrested cell cycle at G2-M phase, which synergized to repress OPM2 cells.

**Figure 4:**
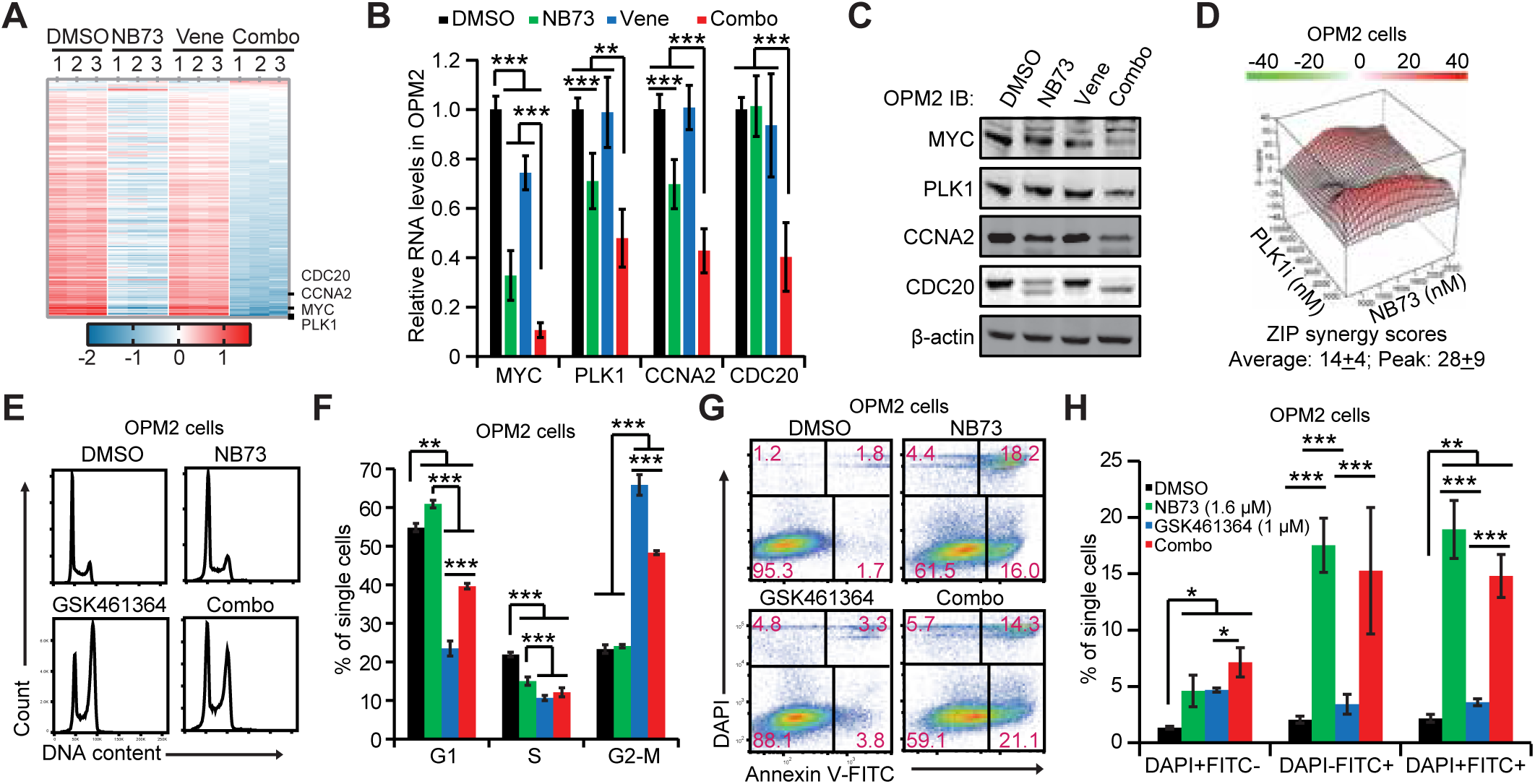
MYC signaling is abrogated by the NB73-Venetoclax combination in OPM2 cells. OPM2 cells were treated with DMSO, NB73, Venetoclax, or their combination for 24 hours for the indicated analysis. (A) A Heatmap depicting all genes in the Hallmark_MYC_Targets_V1 and _V2 gene sets in OPM2 cells. (B) Validation of four down-regulated genes in the MYC pathway through qRT-PCR in OPM2 cells. (C) Representative immunoblotting analysis of these four genes in OPM2 cells. (D) ZIP drug synergy assay of NB73 and a PLK1-specific inhibitor GSK461364 in OPM2 cells. (E) Assessment of cell cycle progression with DAPI-staining in OPM2 cells treated with the indicated drugs for 24 hours. (F) Histogram of cell cycle assays. (G) Assessment of cell apoptosis with Annexin-V binding assay in OPM2 cells treated with the indicated drugs for 24 hours. (H) Histogram of cell apoptosis assays. *p* values were calculated by Student’s t-test with two tails. *: *p*<0.05; **: *p*<0.01; ***: *p*<0.001. n ≥ 3.

Concurrently, we observed similar down-regulation of these four genes by NB73 alone and in combination with Venetoclax at both RNA and protein levels in Delta47 cells (Fig. 5A-B). The average ZIP synergy score of 25.1 demonstrated the synergy of NB73 and GSK461364 in Delta47 cells (Fig. 5C). Surprisingly, GSK461364 not only arrested Delta47 cells at G2-M phase, but also led to sub-G1 population, indicating the cell apoptosis (Fig. 5D-E). The NB73 and GSK461364 combination induced much more cell apoptosis than NB73 or GSK461364 alone (Fig. 5D-E), which was also confirmed by Annexin V-binding assay (Fig. 5F-G). Despite utilizing different mechanisms in OPM2 and Delta47 cells, the combination of NB73 and GSK461364 effectively repressed myeloma cell growth, supporting the repression of MYC pathway by the NB73-Venetoclax pathway.

**Figure 5:**
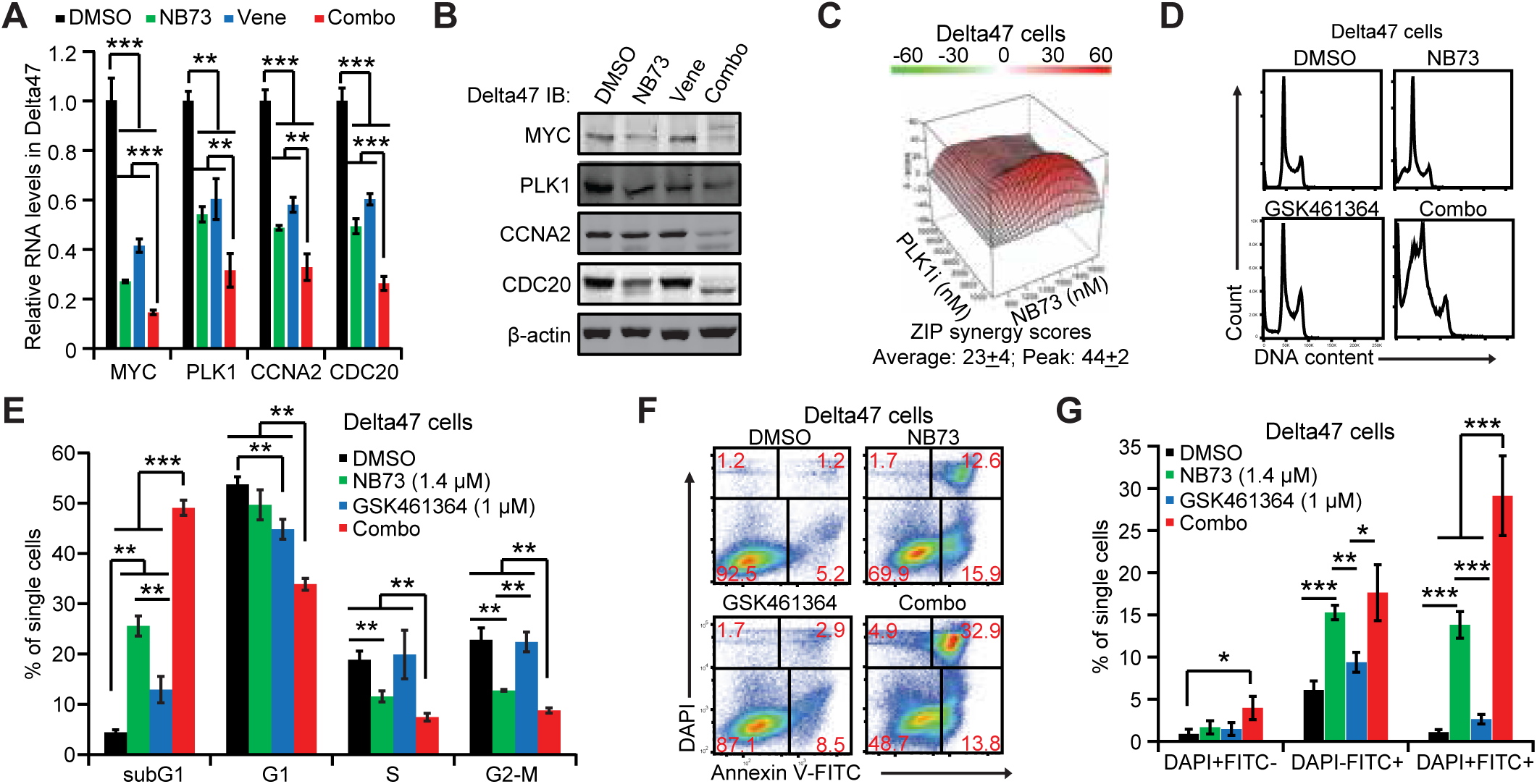
MYC signaling is abrogated by the NB73-Venetoclax combination in Delta47 cells. Delta47 cells were treated with DMSO, NB73, Venetoclax, or their combination for 24 hours for the indicated analysis. (A) Validation of four down-regulated genes in the MYC pathway through qRT-PCR in Delta47 cells. (B) Representative immunoblotting analysis of these four genes in Delta47 cells. (C) ZIP drug synergy assay of NB73 and a PLK1-specific inhibitor GSK461364 in Delta47 cells. (D) Assessment of cell cycle progression with DAPI-staining in Delta47 cells treated with the indicated drugs for 24 hours. (E) Histogram of cell cycle assays. (F) Assessment of cell apoptosis with Annexin-V binding assay in Delta47 cells treated with the indicated drugs for 24 hours. (G) Histogram of cell apoptosis assays. *p* values were calculated by Student’s t-test with two tails. *: *p*<0.05; **: *p*<0.01; ***: *p*<0.001. n ≥ 3.

Furthermore, we investigated the effects of Venetoclax on the MYC pathway in FOXM1-knockout OPM2 and Delta47 cells compared to DMSO control. Under the same experimental conditions as in Figures 4-5, Venetoclax down-regulated the RNA and protein levels of MYC, PLK1, CCNA2, and CDC20 in FOXM1^-/-^ OPM2 cells (supplemental Fig. S2A-B), which was more pronounced compared to Venetoclax-treated FOXM1^+/+^ OPM2 cells (Fig. 4B-C). Similarly, treatment of FOXM1^-/-^ Delta47 cells with Venetoclax also significantly down-regulated the RNA and protein levels of these four genes (supplemental Fig. S2C-D). The difference in the expression of these four genes between FOXM1^-/-^ and FOXM1^+/+^ Delta47 cells was greater than that between FOXM1^-/-^ and FOXM1^+/+^ OPM2 cells. Together, the deletion of FOXM1 alleles enhanced the role of the BCL2 inhibitor Venetoclax in repressing MYC pathway.

### Venetoclax exacerbates the NB73-mediated inhibition of FOXM1 transactivity

NB73 binds to FOXM1 protein for degradation in the proteasome in breast cancer cells [33]. Therefore, we investigated whether NB73 alone and in combination with Venetoclax would regulate FOXM1 expression. As depicted in Figure 6A, NB73 decreased FOXM1 protein levels, while Venetoclax had minimal effect on FOXM1 protein levels in both OPM2 and Delta47 cells. The combination of NB73 and Venetoclax almost completely removed FOXM1 from OPM2 and Delta47 cells, indicating enhanced activity of NB73 in decreasing FOXM1 by Venetoclax (Fig. 6A). Although NB73 reduced the RNA levels of FOXM1 in OPM2 and Delta47 cells, the addition of Venetoclax to NB73-treated cells did not further amplify this inhibition (supplemental Fig. S3), suggesting the need to investigate post-translational regulation of FOXM1 by NB73 alone and in combination with Venetoclax.

**Figure 6:**
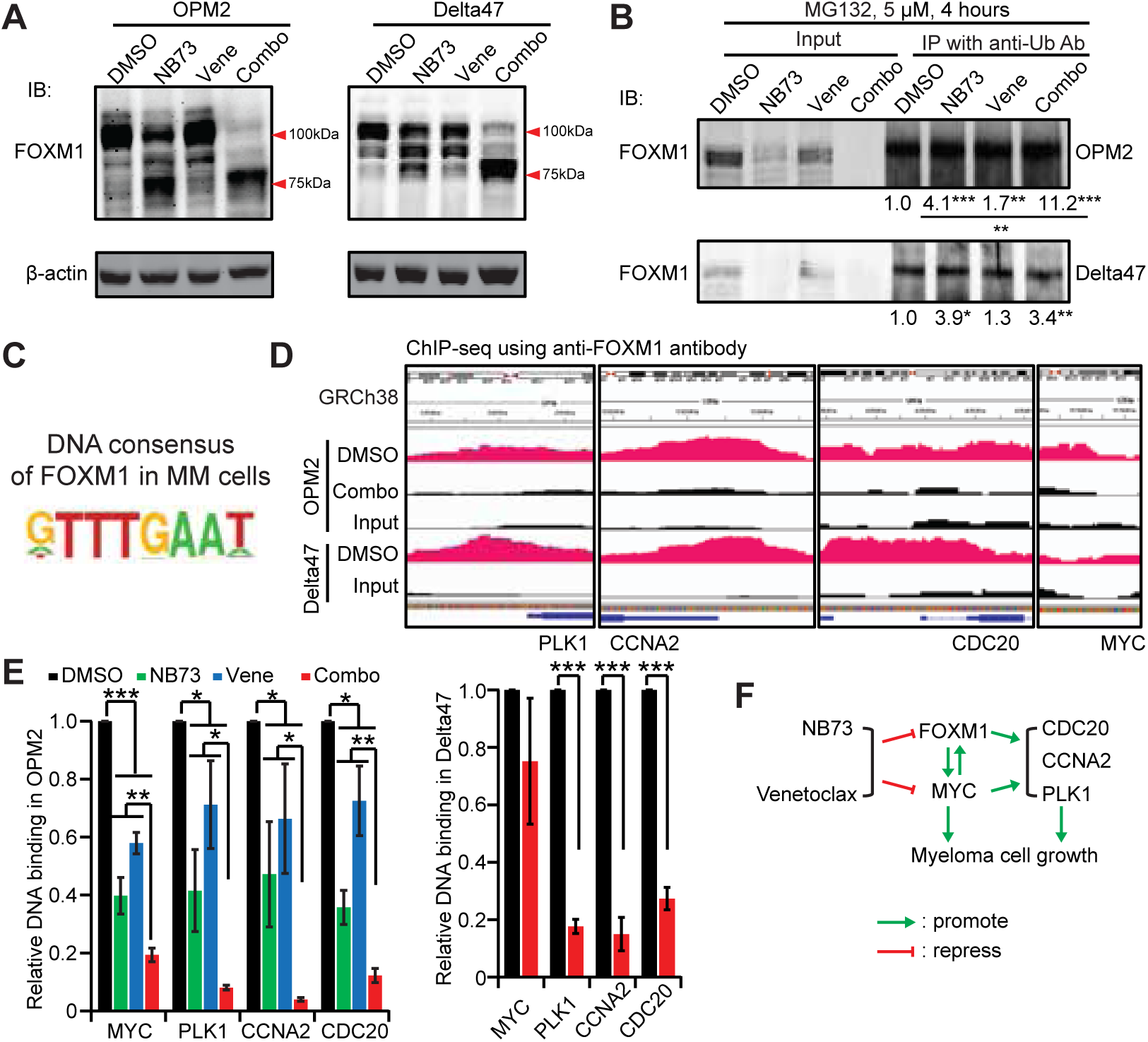
Venetoclax promotes NB73 to decrease FOXM1 and its binding to the promoters of MYC-targeted genes in myeloma cells. Myeloma cells were treated with DMSO, NB73, Venetoclax, or the combination for 24 hours, with or without the proteasome inhibitor MG132. Total cell lysates were then extracted for analysis. (A) Immunoblotting showed representative analysis of FOXM1 levels in OPM2 and Delta47 cells without MG132 addition. (B) Immunoblotting demonstrated FOXM1 levels in total cell lysates and in immunoprecipitates with anti-Ubiquitin antibody in OPM2 and Delta47 cells with MG132 addition. The immunoprecipitate bands were normalized to their input bands, respectively, and the DMSO group was normalized to 1. (C) ChIP-Seq with anti-FOXM1 antibody revealed the DNA consensus of FOXM1 in myeloma cells. (D) Integrative Genomics Viewer images depicted FOXM1-binding sites in the promoters of PLK1, CDC20, CCNA2, and MYC in OPM2 and Delta47 cells treated with DMSO or the NB73-Venetoclax combination, compared to the Input. (E) ChIP-qPCR analyzed FOXM1’s binding to these gene promoters in OPM2 and Delta47 cells using specific primer sets. (F) A proposed molecular model elucidated the NB73-Venetoclax synergy. *p* values were calculated by Student’s t-test with two tails. *: *p*<0.05; **: *p*<0.01; ***: *p*<0.001. n ≥ 3.

Under the same experimental conditions as in Figures 6A, we added the proteasome inhibitor MG132 to the treated cells to block the degradation of ubiquitinated FOXM1 in the proteasome four hours before harvesting the cells. Subsequently, we used an anti-ubiquitin antibody to immunoprecipitate ubiquitinated proteins from the cell lysates and detected ubiquitinated FOXM1 in the immunoprecipitates as well as total FOXM1 in the cell lysates using an anti-FOXM1 antibody. The level of ubiquitinated FOXM1 in the immunoprecipitates was normalized to total FOXM1 in the cell lysates, revealing that NB73 enhanced the ubiquitination of FOXM1 in both OPM2 and Delta47 cells (Fig. 6B). Moreover, the addition of Venetoclax to NB73-treated cells promoted NB73 to enhance FOXM1 ubiquitination in OPM2 cells, resulting in more degradation of FOXM1 (Fig. 6B). However, this Venetoclax-mediated increase was not observed in Delta47 cells, possibly due to the exaggerated degradation of FOXM1 in the NB73 input by the addition of MG132 compared to no MG132 (Fig. 6A-B).

FOXM1 is a master transcription factor in many cancer types [12]. The loss of FOXM1 decreases its binding to the promoters of target genes, thereby losing its regulation of transcriptions (Fig. 4-5). Therefore, we conducted ChIP-seq assays to investigate FOXM1’s binding capabilities to its target genes in OPM2 and Delta47 cells in the presence or absence of the NB73-Venetoclax combination. Compared to DNA input, FOXM1 bound to DNA fragments bearing the “GTTTGAAT” consensus sequence in both OPM2 and Delta47 cells (Fig. 6C). Zooming into the ChIP-seq data, we identified FOXM1-binding sites in the promoters of PLK1, CCNA2, CDC20, and MYC in OPM2 and Delta47 cells (Fig. 6D). qPCR assays using designed primer sets annealing to these binding sites validated FOXM1’s binding to these sites, as well as the decreased binding of FOXM1 in OPM2 cells treated with NB73 (Fig. 6E). The addition of Venetoclax to NB73-treated cells exacerbated the NB73-mediated inhibition of FOXM1’s DNA-binding activities in OPM2 cells (Fig. 6E). Similarly, the combination decreased the FOXM1’s binding to the promoters of CDC20, CCNA2 and PLK1 (Fig. 6E). However, the binding sites of FOXM1 on MYC promoter was not remarkable in Delta47 cells (Fig. 6D), and we did not observe a solid decrease of the FOXM1’s binding to MYC promoter by the NB73-Venetoclax combination after inspecting two possible binding sites with qPCR assay (Fig. 6E & *data not shown*). Together, our studies demonstrate that Venetoclax promotes NB73 to decrease FOXM1 protein levels, abolish FOXM1’s binding to its target genes in MYC pathway, lower the expression of these target genes, repress MYC pathway, and induce myeloma cell apoptosis (Fig. 6F).

## Discussion

### Targeted and combinatorial therapies alleviate drug resistance in MM

The treatment regimens for MM, particularly relapsed/refractory MM and high-risk MM, often involve combinatorial therapies. In this study, our focus has been on developing new FOXM1-targeted MM treatments in combination with known anti-MM drugs. Among these drugs, we discovered the synergy between NB73 and Venetoclax in killing myeloma cells both in vitro (Fig. 1) and in vivo/ex vivo (*in a separate report*). Though Venetoclax is proposed to treat only a subtype of MM with t(11;14) translocation, there are reports on the resistance to Venetoclax in these MM cells, implicating B-cell genes in Venetoclax-sensitive myeloma [8–10]. In contrast, our study has revealed a novel mechanism that FOXM1 plays a critical role in the resistance to Venetoclax in non-t(11;14) MM cells with high FOXM1 expression. In addition to repressing the MYC pathway, NB73 alone or in combination with Venetoclax also decreases BCL2 expression and increases Puma expression in FOXM1^+/+^ OPM2 cells (Fig. 2F), which is consistent with the observations in FOXM1^-/-^ OPM2 cells (supplemental Fig. S1). Therefore, NB73 increases the sensitivity of FOXM1-high non-t(11;14) MM cells to Venetoclax. Our results support the stratification of non-t(11;14) MM cases upon their expression levels of FOXM1 who may take advantage of Venetoclax. Moreover, we are examing the status of FOXM1 in the Venetoclax-resisted t(11;14) MM cell lines [10], and are going to test whether disrupting FOXM1 will also alter their resistance to Venetoclax.

Further studies are indispensable in the ongoing efforts to develop new combinatorial anti-MM therapies. We conducted drug repurposing screens of the NCI AOD X library in FOXM1^-/-^ and FOXM1^+/+^ OPM2 and Delta47 cells, leading to the identification of six new combinations of NB73 with FDA-approved anti-cancer drugs. For example, NB73 synergized with a pan-HDAC inhibitor, Panobinostat that is used in MM [34], in killing OPM2 cells. With the rapid development of computational and AI-guided drug identification techniques, we also utilized the Connectivity Map tool (CMAP) [35] to predict the synergy of NB73 and Thapsigargin, a non-competitive inhibitor of sarcoplasmic/endoplasmic reticulum Ca^2+^-ATPase that stimulates UPR pathway [36]. These two synergies were verified by the ZIP drug synergy test in OPM2 cells (*data not shown*). In contrast, the new-generation FOXM1 inhibitors that were recently published as the first-in-class PROTAC degraders of FOXM1 protein, could serve as an alternative to NB73 [37, 38]. Furthermore, the stable FOXM1^-/-^ cells exhibit a different transcriptomic pattern from the NB73-treated cells (Fig. 2 & supplemental Table-S1), shedding light on conditionally knocking down FOXM1 to develop FOXM1-targeted combinatorial MM treatments.

### Identification of FOXM1-targeted genes in MM

After identifying correlations between FOXM1 and MM stratification and prognosis, we have focused on the molecular mechanisms of FOXM1 in MM progression. We conducted the ChIP-seq assay using the anti-FOXM1 antibody to explore FOXM1 target genes in OPM2 and Delta47 cells (Fig. 6). We proposed the DNA consensus sequence of FOXM1 in myeloma cells as “GTTTGAAT,” which differs from its consensus sequence “TAAACA” suggested in solid tumors [39], or the core consensus sequence “(A/C)AA(C/T)” for the FOX transcription factor family [40]. Not surprisingly, FOXM1 is also well-known for its binding to non-consensus sequences to regulate transcriptions of its target genes [41]. Integrating ChIP-seq data with RNA-seq data led to the recognition of MYC pathway (e.g., MYC, PLK1, CDC20, and CCNA2) as the targets of FOXM1 in MM (Fig. 4-6). These target genes were validated by ChIP-qPCR, qRT-PCR, and immunoblotting in MM cells treated with NB73, Venetoclax, or in combination (Fig. 4-6). Despite previous implications of these four genes as FOXM1 targets [39], we resolved the binding sites of FOXM1 in these genes and validated their transcriptional regulation by FOXM1 using FOXM1 inhibitor NB73 alone and in combination. The synergy between FOXM1-inhibitor NB73 and PLK1-inhibitor GSK461364 also supported the interactions between MYC pathway and FOXM1. One thing remains interesting that FOXM1 may need different partners or cofactors to regulate its target genes in different cells. These partners may be new molecular vulnerabilities in MM which could be targeted therapeutically in precision medicine.

The forkhead DNA-binding domain is structurally similar to linker histone H1, which binds to the entry/exit sites of DNA on the nucleosomal core particle [42–45]. Therefore, FOXM1 may bind to solvent-accessible sites on the nucleosome in repeating intervals, a concept known as “Periodic Binding” [46, 47]. This transcription mode of binding to nucleosomal DNA and recruiting other components to regulate gene expression in transcriptionally inactive and closed chromatin has been known as the “Pioneer Factor” model [47]. In order to investigate this, Transposase-Accessible Chromatin with sequencing (ATAC-seq) in FOXM1-normal and FOXM1-knockout cells will determine if FOXM1 regulates chromatin accessibility across the genome in MM cells. By aligning ATAC-seq, ChIP-seq, and RNA-seq data, the use of multi-omics tools will continue to uncover target genes of FOXM1 in MM progression, identifying additional molecular vulnerabilities of MM for combinatorial anti-MM therapies. In conclusion, we have found that inhibiting FOXM1 increases sensitivity of MM to Venetoclax. The findings reveal the potential benefit of combining a FOXM1 inhibitor for effective treatment of MM.

## Conclusion

Our results have demonstrated that inhibiting FOXM1 increases the sensitivity of non-t(11;14) myeloma cells to BCL2 inhibitor Venetoclax. Considering that non-t(11;14) MM cases comprise 80-85% of all MM cases, our study will shed new light on using Venetoclax in the majority of myeloma cases, which will also increase the response rate of Venetoclax in non-t(11;14) myeloma cases.

## Ethics approval and consent to participate

Not applicable

## Consent for publication

Not applicable

## Availability of data and materials

The materials described in the manuscript, including all relevant raw data, will be freely available to any researcher for non-commercial purposes, without breaching participant confidentiality. The datasets generated during and/or analyzed during the current study are available from the corresponding author on reasonable request.

## Authors’ contributions

Contribution: S.J. and Z.W. designed the FOXM1 project; A.O. instructed and supervised the translational part of the FOXM1 project. Z.W. designed, conducted and analyzed the experiments; S.J. supported the experiments; Y.W. supported the assays completed at UW-Madison; and B.S.K., S.H.K., and J.A.K. supplied the FOXM1 inhibitor NB73 and discussed research findings. Z.W. wrote the manuscript. All authors provided input on and approved the final manuscript.

## Acknowledgements

We extend our sincere gratitude to Dr. Wei Xu at the McArdle Laboratory for Cancer Research, University of Wisconsin-Madison, for graciously hosting and supporting our experiments so that we can use the facilities on the UW-Madison campus. We are also deeply grateful to Dr. Vivian Zhou at the Department of Medicine, Medical College of Wisconsin, for the assistance in materials and methods.

Adam Bissonnette and Luke Moat assisted Z.W. to conduct experiments. Additionally, we would like to express our appreciation to Dr. Todd Kroll, Dr. Abdul Shour, and Dr. Adedayo Onitilo at Marshfield Clinic Health System for their valuable scientific discussions and insights.

