## supplemental figures for "Inhibition of FOXM1 synergizes with BCL2 inhibitor Venetoclax in killing non-t(11;14) multiple myeloma cells via repressing MYC pathway"

### Figure S1, NB73+Venetoclax

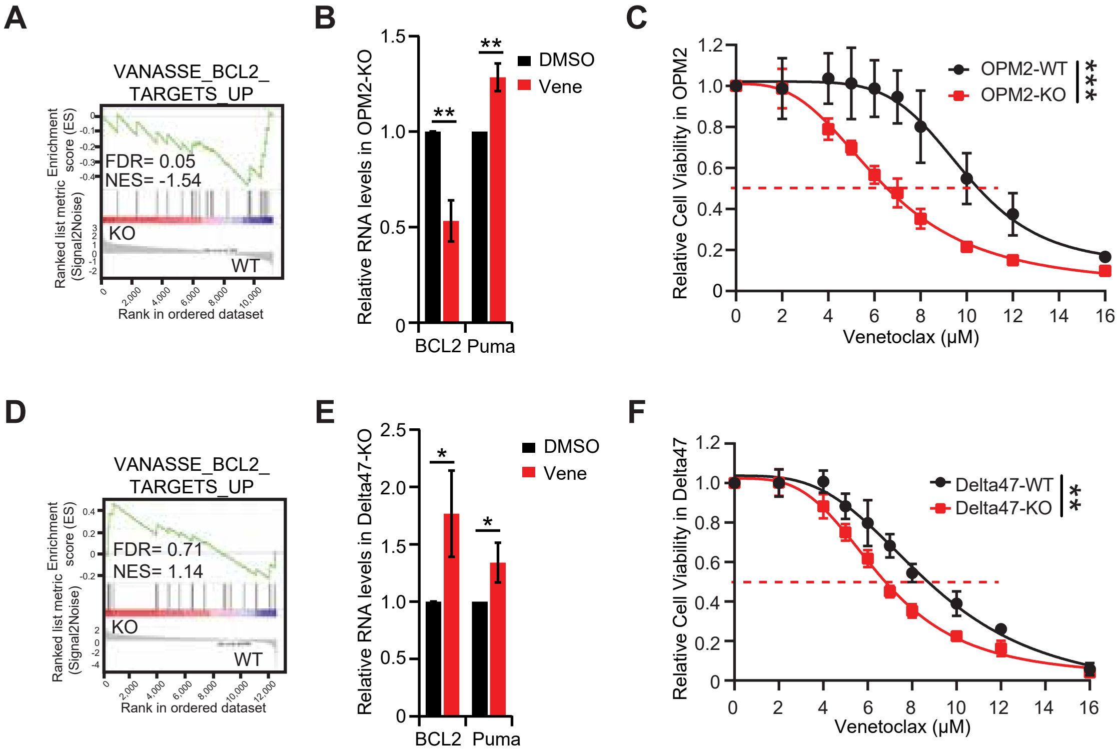

Table-S1: Aligning GSEA studies in NB73 vs DMSO (immediate loss of FOXM1) to these in FOXM1-KO vs FOXM1-normal (stable loss of FOXM1)

| **Enriched in NB73 and FOXM1-normal (opposing)** | **Enriched in DMSO and FOXM1-KO (opposing)** |
|---|---|
| HALLMARK_UNFOLDED_PROTEIN_RESPONSE<br>HALLMARK_APOPTOSIS<br>HALLMARK_INTERFERON_ALPHA_RESPONSE<br>HALLMARK_KRAS_SIGNALING_UP<br>HALLMARK_MTORC1_SIGNALING<br>HALLMARK_TNFA_SIGNALING_VIA_NFKB | HALLMARK_OXIDATIVE_PHOSPHORYLATION<br>HALLMARK_MYC_TARGETS_V1<br>HALLMARK_MYC_TARGETS_V2<br>HALLMARK_E2F_TARGETS<br>HALLMARK_NOTCH_SIGNALING<br>HALLMARK_DNA_REPAIR<br>HALLMARK_G2M_CHECKPOINT<br>HALLMARK_PI3K_AKT_MTOR_SIGNALING<br>HALLMARK_TGF_BETA_SIGNALING<br>HALLMARK_WNT_BETA_CATENIN_SIGNALING<br>HALLMARK_REACTIVE_OXYGEN_SPECIES_PATHWAY |

Pathways listed in the table were from human HALLMARK collection whose FDR values were < 0.25.

#### Figure S2, NB73+Venetoclax

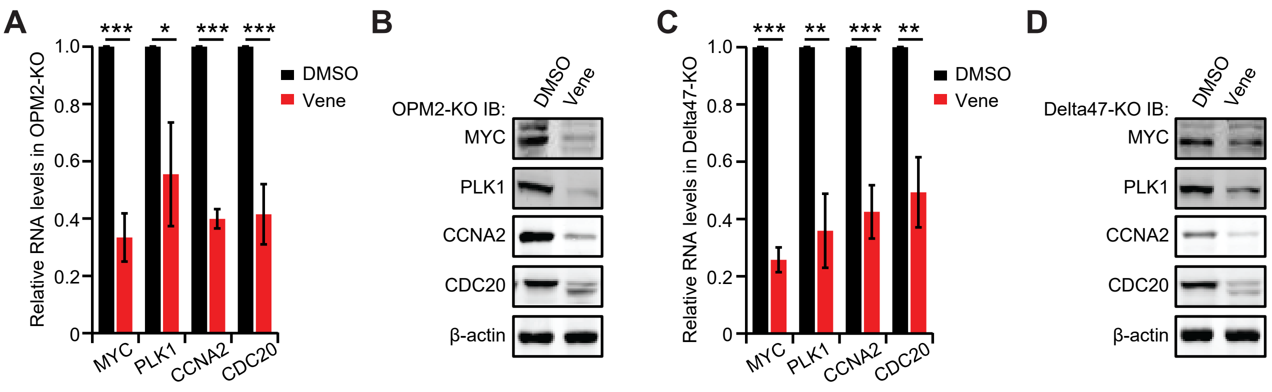

### Figure S3, NB73+Venetoclax

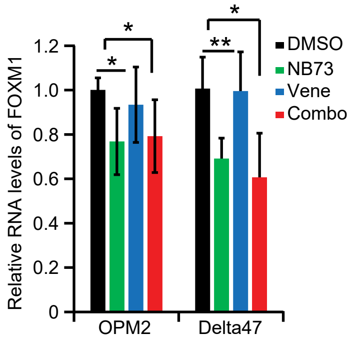
